# Mono-ubiquitination of syntaxin 3 leads to retrieval from the basolateral plasma membrane and facilitates cargo recruitment to exosomes

**DOI:** 10.1101/164996

**Authors:** Adrian J. Giovannone, Elena Reales, Pallavi Bhattaram, Alberto Fraile-Ramos, Thomas Weimbs

## Abstract

Syntaxin 3 (Stx3), a SNARE protein located and functioning at the apical plasma membrane of epithelial cells, is required for epithelial polarity. A fraction of Stx3 is localized to late endosomes / lysosomes though how it traffics there and its function in these organelles is unknown. Here we report that Stx3 undergoes mono - ubiquitination in a conserved polybasic domain. Stx3 present at the basolateral – but not the apical - plasma membrane is rapidly endocytosed, targeted to endosomes, internalized into intraluminal vesicles (ILVs) and excreted in exosomes. A non - ubiquitinatable mutant of Stx3 (Stx3 - 5R) fails to enter this pathway and leads to the inability of the apical exosomal cargo protein GPRC5B to enter the ILV / exosomal pathway. This suggests that ubiquitination of Stx3 leads to removal from the basolateral membrane to achieve apical polarity, that Stx3 plays a role in the recruitment of cargo to exosomes, and that the Stx3 - 5R mutant acts as a dominant - negative inhibitor. Human cytomegalovirus (HCMV) acquires its membrane in an intracellular compartment and we show that Stx3 - 5R strongly reduces the number of excreted infectious viral particles. Altogether these results suggest that Stx3 functions in the transport of specific proteins to apical exosomes and that HCMV exploit this pathway for virion excretion.

## Introduction

SNARE proteins are well recognized as mediators of membrane fusion within the endomembrane system of eukaryotic cells (Rothman, 2014; Sudhof, 2014; Wickner and Schekman, 2008). Syntaxins, a conserved family of SNARE proteins (Weimbs et al., 1997b; Weimbs et al., 1998), are central to all SNARE complexes. At least 16 syntaxins are encoded in the human genome (Hong, 2005), and each localizes to specific membrane domains or organelles in which they carry out specific membrane fusion reactions. The diversity of syntaxins and their cognate SNARE binding partners are thought to contribute to the overall fidelity and specificity of membrane trafficking (Jahn and Scheller, 2006; Rodriguez - Boulan et al., 2005). Others and we have previously reported that Syntaxin 3 (Stx3) localizes to the apical plasma membrane in a wide variety of polarized epithelial cells (Delgrossi et al., 1997; Li et al., 2002; Low et al., 1996; Low et al., 1998; Low et al., 2006; Weimbs et al., 1997a). Apical targeting of Stx3 is governed by an apical targeting signal in its N - terminal domain, and its mutation causes mislocalization of Stx3, improper trafficking of apical membrane proteins, and cell polarity defects (Sharma et al., 2006; ter Beest et al., 2005).

In addition to the apical plasma membrane, a fraction of Stx3 also localizes to late endosomal / lysosomal compartments (Delgrossi et al., 1997; Low et al., 1996). It is unknown how Stx3 traffics to these organelles or what its function there may be. We have previously shown that newly synthesized Stx3 is initially delivered to both the apical and basolateral plasma membrane (Sharma et al., 2006). Because Stx3 is undetectable at the basolateral membrane at steady - state, this suggested that any basolaterally “mis - delivered” Stx3 must be rapidly removed, possibly by endocytosis. Whether and how any syntaxins undergo endocytosis is poorly understood. However, Stx8 interacts with the potassium channel TASK - 1 which facilitates the clathrin - mediated endocytosis of both proteins leading to regulation of TASK - 1 abundance at the plasma membrane (Renigunta et al., 2014). Other syntaxins also interact with ion channels and regulate their activity and / or localization (Bezprozvanny et al., 1995; Geng et al., 2008; Leung et al., 2007; Singer - Lahat et al., 2008) which has led to the idea that syntaxins have additional functions unrelated to membrane fusion.

Curiously, Stx3 has been identified among proteins found in apically secreted exosomes but not basolateral exosomes (van Niel et al., 2001). In vivo, Stx3 is also found in urinary exosomes, presumably due to apical secretion from tubule epithelial cells (Gonzales et al., 2009). Exosomes are small vesicles that are secreted by the fusion of intracellular multivesicular bodies (MVBs) with the plasma membrane (Lakkaraju and Rodriguez - Boulan, 2008; Meckes and Raab - Traub, 2011). MVBs, in turn, are a class of late endosomes containing intraluminal vesicles (ILVs) which are formed by the invagination of the limiting membrane of the endosome. MVBs either fuse with lysosomes for the degradation of their membranous contents or with the plasma membrane for the secretion of exosomes. Conjugation of membrane proteins with ubiquitin can serve as a signal that directs these membrane proteins into the MVB pathway (Huang et al., 2006; Kamsteeg et al., 2006; Marchese et al., 2008; Stringer and Piper, 2011). In particular, monoubiquitination has been identified as an endocytosis signal that directs plasma membrane proteins to the endocytic pathway (Hicke, 2001).

Human cytomegalovirus (HCMV) is a member of the enveloped herpesvirus family, and a widespread human pathogen that causes asymptomatic infections that often become latent. However, in immunocompromised persons, HCMV infections can be life - threatening, and congenital infection can lead to significant developmental defects (Britt and Mach, 1996; Fowler et al., 1992). HCMV acquires its final envelope in intracellular membranes prior to its secretion though the mechanisms underlying these processes are poorly understood. We have previously shown that HCMV strongly induces the expression of Stx3 and that Stx3 is required for the production of infectious viral particles through a mechanism that likely involves late endosomes/lysosomes (Cepeda and Fraile - Ramos, 2011). How Stx3 may be involved in HCMV virion production and secretion is unknown. However, according to some models, HCMV virions bud into intracellular membrane organelles which may subsequently fuse with the plasma membrane in a mechanism that at least superficially resembles the mechanism of secretion of exosomes from MVBs (Alenquer and Amorim, 2015; Cepeda et al., 2010; Fraile - Ramos et al., 2010).

We report here that Stx3 can undergo mono - ubiquitination on lysines in its juxtamembrane region. Stx3 is rapidly endocytosed from the basolateral domain of polarized epithelial cells but not from the apical domain. A ubiquitination - deficient mutant of Stx3 (Stx3 - 5R) exhibits delayed basolateral endocytosis, exclusion from the endosomal pathway and diminished secretion in apical exosomes. Expression of Stx3 - 5R acts in a dominant - negative fashion and decreases the exosomal secretion of GPRC5B, a known apical exosome cargo. Finally, expression of Stx3 - 5R strongly inhibits the number of infectious HCMV secreted virions. Altogether, these results suggest that Stx3 has a second function in addition to its established role in apical membrane fusion. Stx3 is ubiquitinated and actively transported from the basolateral membrane to the endosomal/MVB pathway and excreted apically in exosomes. Our results suggest that Stx3 is not only a passive cargo in this pathway but facilitates the efficient trafficking of other cargoes into apical exosomes. In addition, our results also suggest that HCMV exploits this cellular apical exosomal pathway to accomplish the efficient release of viral particles.

## Results

### Stx3 undergoes mono - ubiquitination

While the bulk of Stx3 resides on the apical plasma membrane in polarized epithelial cells, a portion of Stx3 also localizes to a late endosomal population (Delgrossi et al., 1997; Low et al., 1996). We hypothesized that ubiquitination may be involved in the targeting of endosomal Stx3. To test whether Stx3 can undergo ubiquitination, endogenous Stx3 was immunoprecipitated from polarized Caco2 intestinal epithelial cells and probed for ubiquitin by immunoblotting. A specific, ubiquitin - positive band was detected migrating at a position ∼9 kDa larger than Stx3 (Fig. 1A) suggesting that a fraction of Stx3 undergoes mono - ubiquitination. To confirm this finding, myc - tagged ubiquitin and untagged Stx3 were co - expressed in HEK293T cells followed by immunoprecipitation. Mono - ubiquitinated Stx3 can be detected as a band with an increased molecular weight of ∼9 kDa (Fig. 1B). Using MDCK cells stably expressing either C - terminally myc - tagged Stx3 or Stx4 in a doxycycline - inducible manner (Low et al., 2006), a similar ∼9 kDa shifted, ubiquitin - positive band appears for Stx3 but not Stx4 suggesting that ubiquitination is not a universal modification of all syntaxins (Fig. 1C). Treatment with the protease inhibitor ALLN leads to a strong increase in the level of mono - ubiquitinated Stx3 (Fig. 1D). The effect of ALLN suggests that the ubiquitinated species is either quickly degraded or de - ubiquitinated in the cell. ALLN has previously been suggested to inhibit deubiquitinating enzymes (Wojcikiewicz et al., 2003). Altogether, these data indicate that a short - lived population of Stx3 can undergo mono - ubiquitination in multiple cell lines and under various experimental conditions.

**Figure 1.**
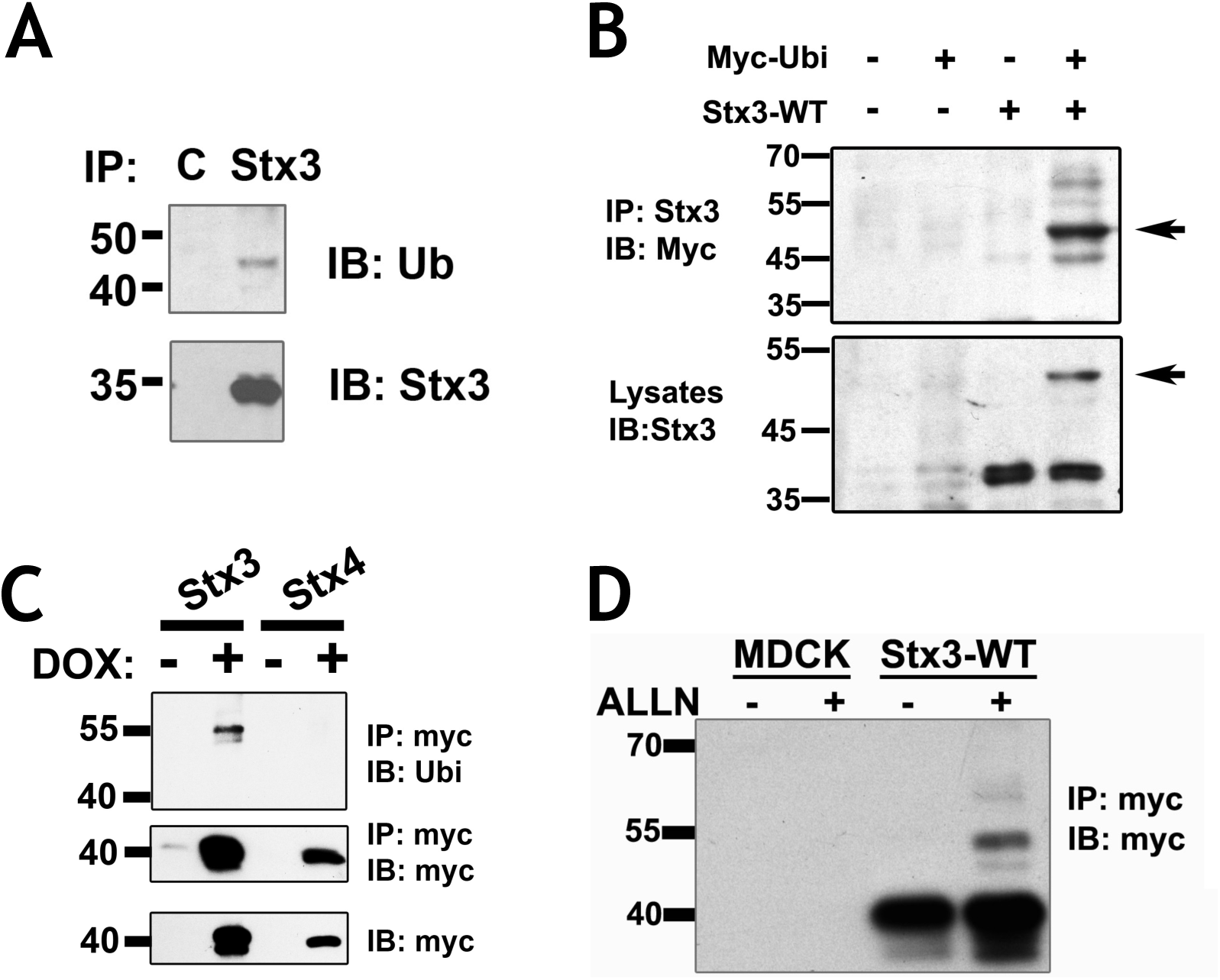
Stx3 is ubiquitinated. (A) Endogenous Stx3 was immunoprecipitated (IP) from Caco2 cells and immunoblotted (IB) with an antibody to ubiquitin (Ub) and Stx3. Nonspecific rabbit IgG antibody was used for control (lane C). (B) HEK293T cells were transfected with untagged Stx3 and myc - ubiquitin constructs. Lysates were subjected to IP with an anti - Stx3 antibody and analyzed by IB with anti - myc antibody. Cell lysates used as input for the IP were blotted with anti - Stx3 antibody. The ubiquitinated Stx3 band identified by both anti - myc and anti - Stx3 antibodies is indicated by arrow. (C) Doxycycline inducible MDCK cells expressing Stx3 or Stx4, both myc - tagged, were lysed, immunoprecipitated with anti - myc antibodies, and analyzed by immunoblot with antibodies to myc or Ubi. Bottom panel: total lysates used for the IP were blotted with anti - myc antibody. (D) Untransfected MDCK cells or MDCK cells stably expressing double myc - tagged Stx3 were treated with 10μM ALLN (+ ALLN) or without (- ALLN) for 16h. Lysates were subjected to IP with anti - myc antibody and analyzed by IB with anti - myc antibody.

### Ubiquitination occurs in the juxtamembrane region of Stx3

Deletion mutants were used to identify the ubiquitination site(s) in Stx3 (Fig. 2A). Deletion of the transmembrane domain prevents ubiquitination suggesting that membrane anchorage is required (Fig. 2B). Constructs containing either the N - or C - terminal halves, respectively, of the cytoplasmic domain of Stx3 in addition to the transmembrane domain could both be ubiquitinated (Fig. 2). The only lysine residues in common between these two constructs are the two lysines immediately preceding the transmembrane domain which are part of a basic motif containing six lysine residues (Fig. 2A). Similar basic motifs are present in most other SNARE proteins that contain C - terminal transmembrane anchors (Weimbs et al., 1998) but the primary amino acid sequences of these motifs are not well conserved between different syntaxins (Fig. 2C). To investigate possible ubiquitination in this basic motif of Stx3, lysines were progressively replaced with arginines in order to prevent ubiquitination without disrupting the positive charges (Fig. 2D). Mutation of all six lysines (6R mutant) or of the most C - terminal five lysines (5R mutant) completely prevented ubiquitination (Fig. 2E). Interestingly, the 2R, 3R, and 4R mutants did not prevent ubiquitination. Given the positions of these mutations we conclude that any of the C - terminal five lysine residues can be subject to ubiquitination. This suggests that ubiquitination of this motif is region - specific but not strictly sequence specific.

**Figure 2.**
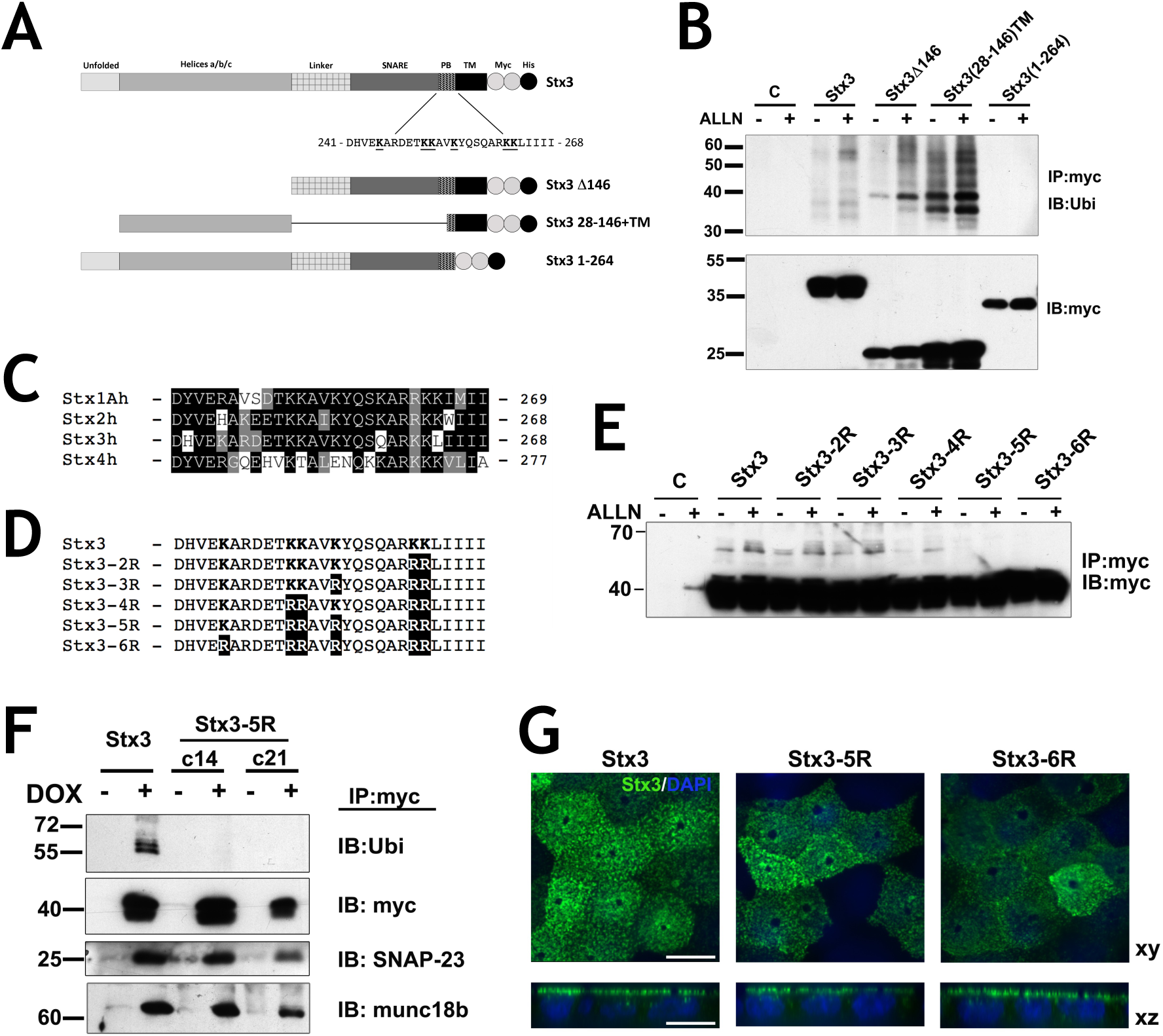
Ubiquitination occurs in the juxtamembrane region of Stx3. (A) Schematic representation of Stx3 wild - type and mutant constructs used for expression. Residues in the polybasic juxtamembrane region (PB) of wild - type Stx3 are shown. Two myc epitope tags (white circles) and one His6 tag (black circle) were added to the COOH termini; PB, polybasic; TM, transmembrane domain. (B) HEK293T cells were transiently transfected with the indicated constructs, plated for 24h and incubated with or without ALLN. Lysates were subjected to IP with anti - myc antibody and analyzed by IB with anti - Ubi. Bottom: Total lysates used for IP blotted with anti - myc antibody. Cells without construct transfection (lane C) serve as a negative control for the IP. (C) Sequence alignment of the juxtamembrane region of human plasma membrane syntaxins 1A, 2, 3 and 4, SwissProt accession numbers Q16623, P32856, Q13277, Q12846 respectively. (D) Stx3 lysine mutant constructs used in this study (region 241 - 268 is shown). Lysine residues mutated to arginine are in bold letters. (E) Anti - myc Western Blot of HEK293T treated with and without ALLN transiently expressing lysine mutants subjected to anti - myc IP. (F) Two different stable clones of DOX inducible MDCK cells stably transfected for Stx3 - 5R, c14 and c21, were subjected to IP with anti - myc antibody and IB with antibodies against Ubi, SNAP - 23, and munc18b. (G) Immunocytochemistry of DOX - induced Stx3, Stx3 - 5R, or Stx3 - 6R (green) expresses in polarized MDCK cells cultured on Transwell filters. Scale bar: 10μm.

We chose the “5R” mutant as a non - ubiquitinatable Stx3 for further study and stably expressed this mutant in MDCK cells in a doxycycline - inducible manner. Stx3 - 5R still binds efficiently to the endogenous SNARE binding partner SNAP - 23 and the SNARE regulator munc18b similar to wild - type Stx3 (Fig. 2F) suggesting that the mutations in Stx3 - 5R do not cause detrimental structural defects that disrupt normal binding interactions. The steady - state localization of Stx3 - 5R and Stx3 - 6R at the apical plasma membrane mirrors wild - type localization (Fig. 2G).

### Ubiquitination directs basolateral Stx3 to the endocytic pathway and influences cargo sorting

Previously, we have reported that, in addition to the apical plasma membrane, a fraction of Stx3 localizes to late endosomes / lysosomes (Low et al., 1996). To test if this localization is ubiquitination - dependent, we investigated the intracellular localization of wild - type Stx3 or Stx3 - 5R by immunofluorescence microscopy in MDCK cells. As expected, wild - type Stx3 extensively colocalizes with the late endosomal / lysosomal marker LAMP - 2. In contrast, Stx3 - 5R exhibits very little colocalization with LAMP - 2 (Fig. 3A) suggesting that ubiquitination may be required for Stx3 targeting into the late endosomal / lysosomal pathway.

**Figure 3.**
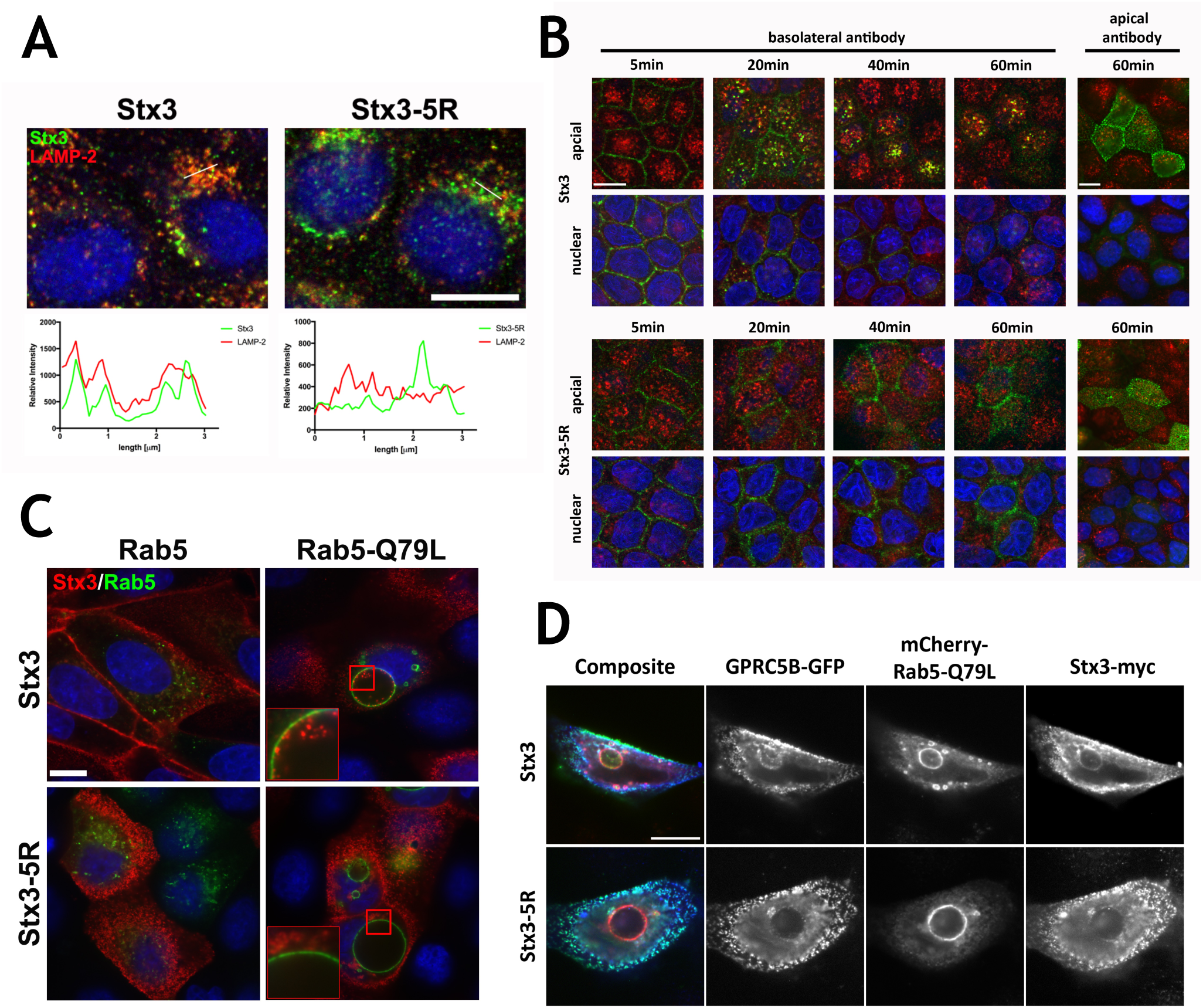
Ubiquitination directs basolateral Stx3 to the endocytic pathway and influences cargo sorting to endosomes. (A) Immunofluorescence microscopy showing reduced colocalization of Stx3 - 5R with LAMP - 2 compared to wild - type Stx3. Cells were cultured on coverslips, treated with DOX for 16h and stained with Stx3 (green) and LAMP - 2 (red) antibodies. Intensity of pixels along white lines in micrographs for each channel plotted in graphs under micrographs. Scale bar: 10μm. (B) Wild - type Stx3 or Stx3 - 5R (c14) expression was induced with DOX for 16h in stably transfected MDCK cells grown at confluence for 4 days on Transwell filters. For endocytosis assay anti - myc 9E10 ascites was added to apical or basolateral media and incubated at 4°C before being washed off and incubated at 37°C for indicated times. Cells were fixed and stained for Stx3 (myc), green; M6PR, red; and nuclei (DAPI), blue. Scale bar: 10 μm. (C) MDCK cells stably expressing Stx3 or Stx3 - 5R were grown on coverslips and transiently transfected with GFP - Rab5 or GFP - Rab5Q79L. Twenty - four hours after transfection, Stx3 or Stx3 - 5R expression was induced with DOX for 16h. Cells were fixed and processed for immunocytochemistry. Stx3 (myc), red; Rab5, green; and nuclei (DAPI), blue. Scale bar: 10μm. (D) MDCK cells were grown on coverslips and transiently transfected with mCherry - Rab5 - Q79L, GFP - GPRC5B, and either Stx3 or Stx3 - 5R. Thirty - six hours after transfection cells were fixed and processed for immunocytochemistry. Stx3 (myc), blue; Rab5, red; and GPRC5B, green. Scale bar: 10μm.

To investigate this novel role for ubiquitination of Stx3, we took advantage of the extracellular myc - epitope tag (Fig. 2A) to follow the endocytosis of Stx3 from either the basolateral or apical plasma membrane. Polarized MDCK cells cultured on Transwell filters were incubated with anti - myc antibody either in the media compartment in contact with the apical or the basolateral domain. After antibody addition, cells were washed, and incubated at the indicated time points. Any Stx3 that had been tagged with the antibody on either cell surface was visualized by immunofluorescence microscopy (Fig. 3B). Surprisingly, despite the fact that wild - type Stx3 is undetectable at the basolateral membrane at steady - state (Fig. 2G), after antibody addition into the basolateral chamber a strong signal of antibody - tagged Stx3 is observed on the basolateral membrane at 5 minutes (Fig. 3B). Within 20 minutes, antibody - tagged Stx3 moves to intracellular vesicles that co - stain with the M6PR (mannose 6 - phosphate receptor), a marker of the late endosomal / lysosomal pathway. Most of the basolaterally internalized Stx3 signal remains in M6PR - positive organelles after 60 minutes but a fraction appears to be able to reach the apical plasma membrane by that time. In contrast, when the myc - antibody is added to the apical chamber antibody - tagged wild - type Stx3 remains at the apical membrane with no evidence of internalization after 60 minutes (Fig. 3B). We conclude that a significant fraction of wild - type Stx3 is targeted to the basolateral domain from which it is rapidly removed by endocytosis followed by targeting to the late endosomal / lysosomal pathway. In contrast, the fraction of Stx3 that has reached the apical membrane is stable at this location and does not undergo endocytosis. Therefore, apical polarity of Stx3 is achieved, at least in part, by selective removal from the “incorrect” plasma membrane domain.

The ubiquitination - deficient Stx3 - 5R mutant is still efficiently tagged by the myc - antibody at both the basolateral and apical membranes. In contrast to wild - type Stx3, however, Stx3 - 5R does not undergo rapid endocytosis from the basolateral membrane, and does not exhibit targeting to M6PR - positive organelles. Eventually, by 60 minutes, a fraction of Stx3 - 5R reaches the apical membrane. We conclude that the inability to ubiquitinate Stx3 leads to a defect in efficient basolateral endocytosis and targeting to the late endosomal / lysosomal pathway.

To further explore the internalization and endosomal trafficking of Stx3 we utilized the GTPase - deficient Q79L mutant of Rab5 (Barbieri et al., 1996). Proteins internalized via clathrin - mediated endocytosis accumulate in early endosomes in the presence of this mutant and these endosomes dramatically enlarge in size (Gong et al., 2007; Olkkonen and Stenmark, 1997; Somsel Rodman and Wandinger - Ness, 2000; Zerial and McBride, 2001). Stx3 is efficiently targeted to Rab5 - Q79L - positive enlarged endosomes and localizes both at the limiting membrane and prominently on intraluminal vesicles (ILVs) of these endosomes (Fig. 3C). This result suggests that Stx3 is internalized to ILVs. In contrast, Stx3 - 5R is absent from both the limiting membrane and ILVs of Rab5 - Q79L - positive enlarged endosomes suggesting again that ubiquitination is required for endocytosis of Stx3 and trafficking to endosomes (Fig. 3C).

The orphan G - protein coupled receptor GPRC5B is excreted with apical exosomes in MDCK cells and has been shown to localize to ILVs in enlarged Rab5 - Q79L - positive early endosomes (Kwon et al., 2014). Therefore, we investigated GPRC5B as an established cargo protein that traffics via endosomal ILVs en route the apical exosomal pathway in MDCK cells. mCherry - tagged Rab5 - Q79L, GFP - tagged GPRC5B and Stx3 or Stx3 - 5R were co - expressed in MDCK cells. As anticipated, GPRC5B and Stx3 localize to the lumens of enlarged early endosomes (Fig. 3D). However, when Stx3 - 5R is expressed, neither Stx3 - 5R itself nor GPRC5B are found in the endosomal lumen. This result suggests the possibility that Stx3 not only traffics to ILVs in the endosomal pathway but may also play a role in facilitating the trafficking of other proteins in this pathway. Overexpression of the ubiquitination - defective Stx3 - 5R mutant may therefore act as a dominant - negative inhibitor of this pathway.

### Ubiquitination directs Stx3 and cargo to exosomes

Stx3 and its binding partner munc18b have been identified in apically released exosomes from intestinal epithelial cells (van Niel et al., 2001) and in a proteomics screen of urinary exosomes (Gonzales et al., 2009). We confirmed the presence of Stx3 and munc18b in human urinary exosomes (Fig. 4A). Besides the relatively abundant signal for Stx3 in urinary exosomes, Stx2 could also be detected (Fig. 4B), but not the basolateral membrane - specific Stx4 (Low et al., 1996) or the neuronal Stx1A (Fig. 4B). Next, we isolated exosomes secreted from Caco - 2 cells and found endogenously expressed Stx3 to be present (Fig. 4C). These results confirm that endogenous Stx3 is secreted in exosomes both *in vivo* and *in vitro*.

**Figure 4.**
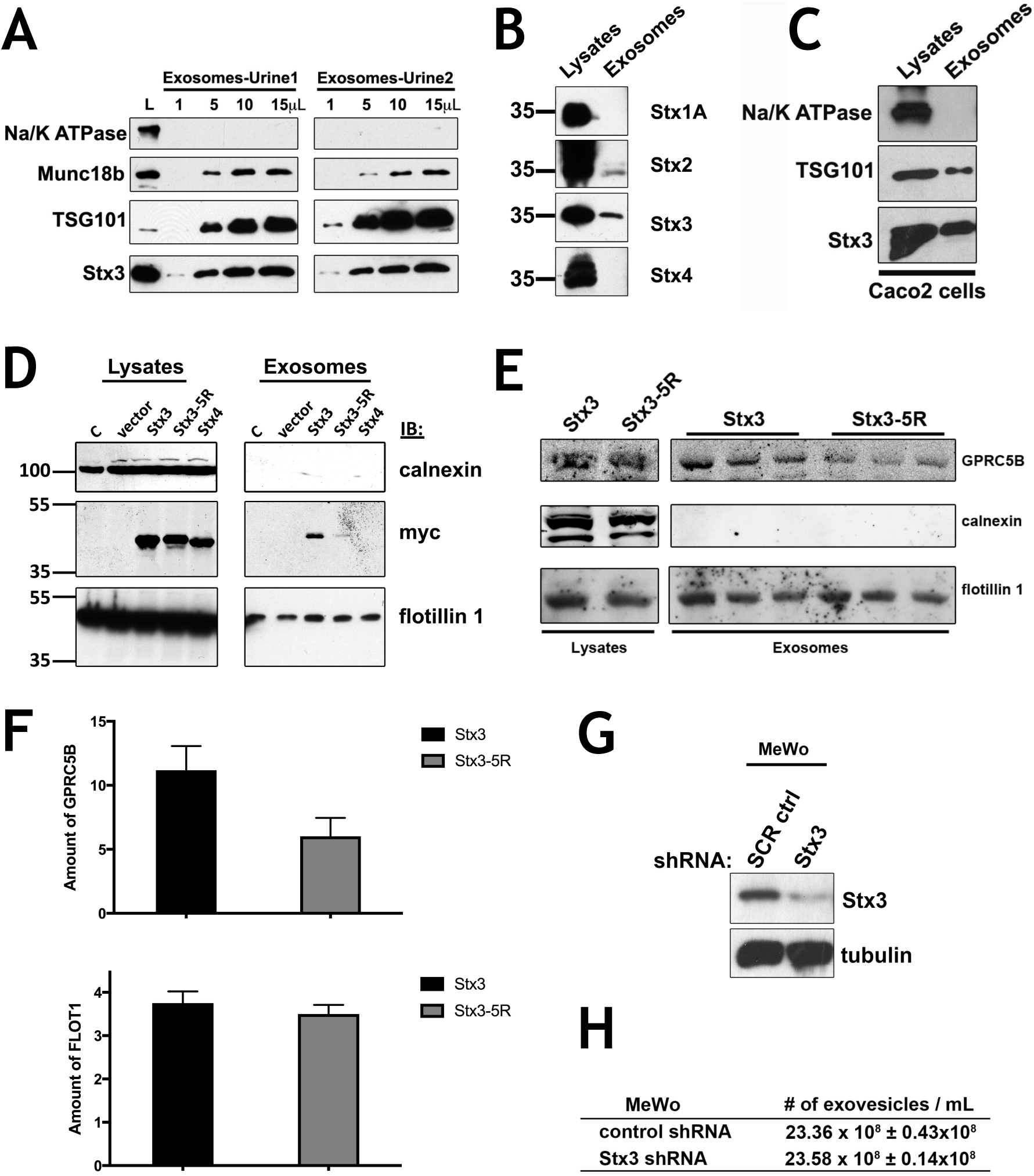
Ubiquitination directs Stx3 and cargo to exosomes. (A) Exosomes from 60mL of human urine (from two volunteers 1 and 2) were isolated by differential ultracentrifugation. Indicated volume amount from a total volume of 50μL were analyzed by immunoblotting for TSG101 (exosome positive marker), Na/K ATPase (exosome negative marker) (Simons and Raposo, 2009; Thery et al., 2009), Stx3, and munc18b. Total lysates from Caco - 2 cells were used as antibody control (L) (B) Human urine exosomes were analyzed for the indicated plasma membrane syntaxins. Lysates from MDCK cells with stable expression for the respective syntaxins were used as positive controls. (C) Exosomes from Caco - 2 conditioned media were isolated and analyzed as above. (D) HEK293T cells were transiently transfected with Stx3, Stx3 - 5R, and Stx4 (all myc tagged) plasmids. Lane C represents cells without transfection. Forty - eight hours after transfection exosomes from the media and cell lysates were analyzed by immunoblotting with anti - myc, anti - calnexin (exosome negative marker) and anti - flotillin - 1 (exosome positive marker). As a specificity control, Stx4 was included which was not appreciably secreted in exosomes. (E) DOX induced MDCK expressing Stx3 or Stx3 - 5R were transiently transfected with GFP - GPRC5B. Exosomes were purified from media after 48h and subjected to immunoblotting. Performed in triplicate. (F) Quantification of relative intensity of GPRC5B bands in Stx3 or Stx3 - 5R exosome samples from (E). Error bars represent SEM of experiment performed in triplicate. *P< 0.05, Student’s unpaired t - test. (G) Immunoblot of MeWo cells transduced with lentivirus delivering shRNA #304, targeting Stx3, or shRNA scrambled control. (H) Table showing concentration of exovesicles in media collected from cells in panel (G).

To determine if ubiquitination is necessary for exosomal secretion of Stx3, we expressed myc - tagged Stx3 or Stx3 - 5R in HEK293T cells and isolated exosomes from the cell culture medium. While wild - type Stx3 is readily detectable in exosomes, only trace amounts of Stx3 - 5R were found (Fig. 4D) suggesting that ubiquitination is required to direct Stx3 to the endosomal pathway leading to secretion in exosomes.

Since our above data suggested that Stx3 may play a role in facilitating the trafficking of GPRC5B into the endosomal pathway, we next investigated whether the exosomal secretion of GPRC5B is affected by Stx3 - 5R. GPRC5B was expressed in MDCK cells stably expressing either Stx3 or Stx3 - 5R and the secreted exosomes were isolated. Secretion of the general exosomal marker flotillin was unaffected by expression of Stx3 or Stx3 - 5R (Fig. 4E). In contrast, the exosomal secretion of GPRC5B was reduced by approximately 50% in cells expressing Stx3 - 5R (Fig. 4F). This result suggests that Stx3 may facilitate the exosomal trafficking of specific cargo proteins as opposed to the secretion of exosomes *per se*. To further test whether Stx3 is required for the overall ability of cells to secrete exosomes we knocked down the expression of endogenous Stx3 by stable shRNA expression in MeWo cells (Fig. 4G) and quantified the number of secreted exosomal particles by Nanoparticle Tracking Analysis (Carr, 2007). No substantial difference was observed (Fig. 4H).

Altogether, these results suggest that Stx3 is not required for the biogenesis and secretion of exosomes but likely plays a role in the targeting of specific proteins into the exosomal pathway.

### Ubiquitination of Stx3 is required for HCMV secretion

We have previously shown that HCMV infection induces a strong increase in the expression of endogenous Stx3, and that Stx3 can be detected in secreted virions (Cepeda and Fraile - Ramos, 2011). Knockdown of Stx3 expression inhibits the production and secretion of HCMV virions (Cepeda and Fraile - Ramos, 2011), a processes that also involves Rab27a (Fraile - Ramos et al., 2010), a small GTPase required for exosome excretion (Ostrowski et al., 2010). These findings suggest that Stx3 is required for a key step in the HCMV life cycle and we hypothesized that its role may depend on its ability to be ubiquitinated and target to the endosomal / exosomal pathway. To test this, the expression of endogenous Stx3 in human foreskin fibroblast (BJ1) cells was knocked - down by shRNA (Fig. 5A) and then “rescued” by lentiviral expression of Stx3 or Stx3 - 5R cDNA constructs resistant to the shRNA (Fig. 5A). When these cells were infected with a recombinant strain of HCMV containing a GFP reporter under an HCMV early promoter (McSharry et al., 2001) and the number of infected cells was determined by GFP expression, we did not observe differences between the cell lines (Fig. 5B). Knock - down of endogenous Stx3 expression caused a strong reduction of the amount of infectious particles released into the supernatant (Fig. 5C) consistent with previous results (Cepeda and Fraile - Ramos, 2011). This effect was completely rescued by re - expression of wild - type Stx3 but not by re - expression of Stx3 - 5R (Fig. 5C). Moreover, exogenous expression of Stx3 - 5R alone – without knocking - down endogenous Stx3 expression - strongly reduced the number of infectious particles released (Fig. 5C) indicating again that this mutant acts as a dominant - negative inhibitor. Altogether, these results indicate that ubiquitination of Stx3 plays a key role in the life cycle of HCMV and suggest that HCMV uses a pathway similar to the exosomal pathway for its secretion.

**Figure 5.**
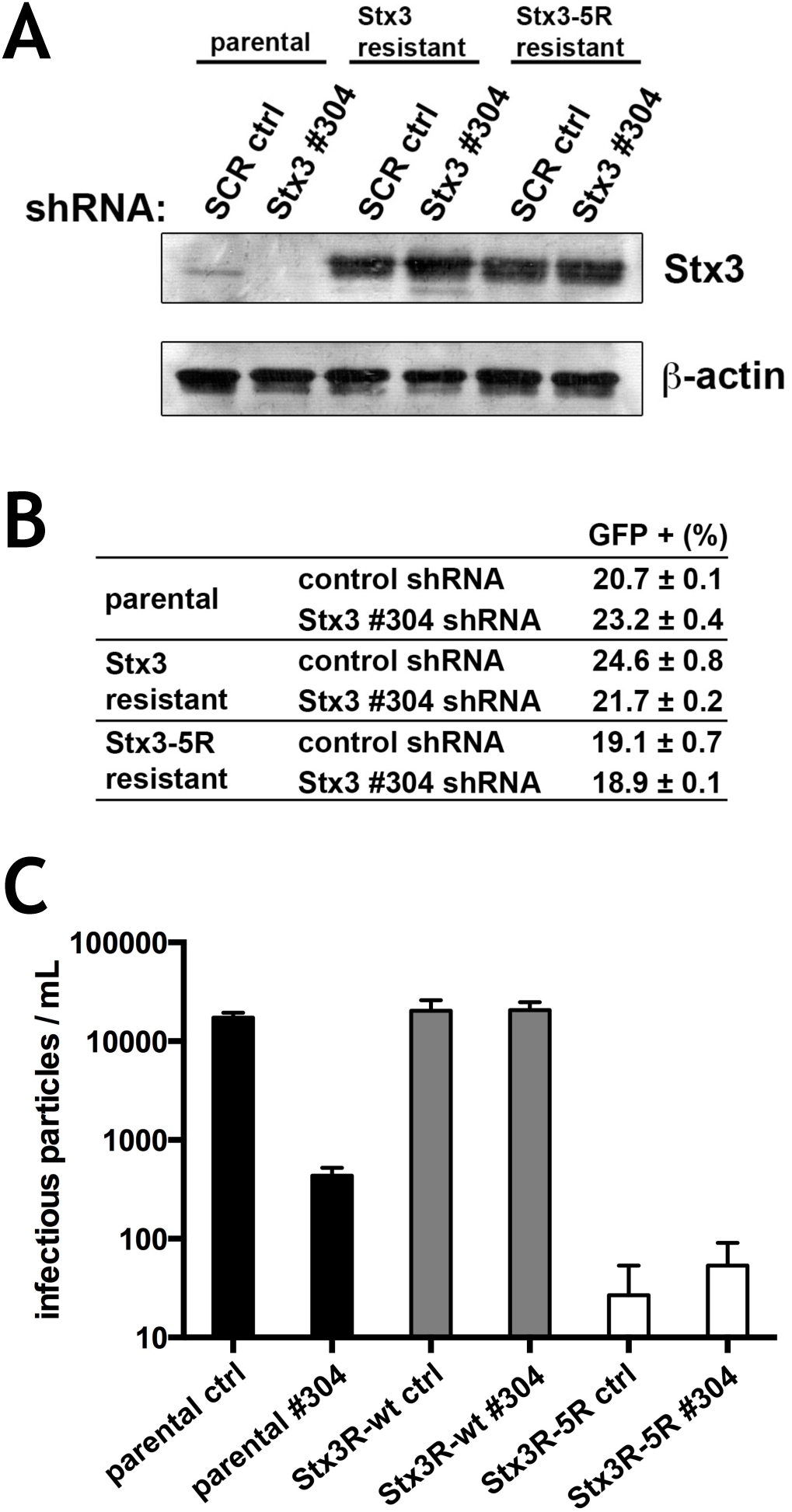
Ubiquitination of Stx3 is required for HCMV secretion. BJ1 cells expressing shRNA - resistant - STX3 or shRNA - resistant - Stx3 - 5R constructs were transduced with scrambled control and STX3 #304 shRNAs and infected with RecCMV at 0.3 moi. (A) Immunoblot analysis of these cells at 4 days post infection with antibodies against STX3 and actin. (B) Flow cytometry analysis for GFP expression. (C) At 5 dpi, supernatants were harvested and the number of extracellular infectious viruses was determined on fresh BJ1 cells. Data are means plus standard deviations of experiment performed in quadruplicate.

## Discussion

This study illustrates a novel trafficking pathway for Stx3 and suggest a function of Stx3 that may be independent of its function in membrane fusion. Previous studies have shown that Stx3 localizes to the apical plasma membranes of various types of polarized epithelial cells and that it plays a role in apical membrane trafficking, presumably by mediating fusion of transport vesicles that reach the apical plasma membrane (Breuza et al., 2000; Delgrossi et al., 1997; Fujita et al., 1998; Li et al., 2002; Naren et al., 2000; Riento et al., 1998). However, Stx3 has also been detected on intracellular membrane compartments including late endosomes / lysosomes (Delgrossi et al., 1997; Low et al., 1996), secretory granules in pancreatic acinar cells (Gaisano et al., 1996), mast cells (Guo et al., 1998; Hibi et al., 2000) and parotid acinar cells (Castle et al., 2002), tubulovesicles containing the H,K - ATPase in gastric parietal cells (Peng et al., 1997), the phagosomal membrane of macrophages (Hackam et al., 1996), and on melanosomes in melanocytes (Yatsu et al., 2013). It is unclear whether Stx3 functions in these intracellular organelles as a membrane fusion protein or whether it may have a separate function.

In MDCK cells, newly synthesized Stx3 reaches both the apical and basolateral plasma membranes on direct routes after passage through the Golgi in a relatively non - polarized fashion (Sharma et al., 2006) and it remained unclear how apical polarity is eventually achieved. In addition, it has remained unknown how Stx3 may be targeted to any of the intracellular organelles mentioned above. Our results now suggest that Stx3 is targeted to the endosomal / lysosomal pathway via endocytosis from the basolateral plasma membrane. Our finding that myc - tagged Stx3 can be efficiently detected by binding to the myc antibody at the basolateral membrane (Fig. 3B) is consistent with our previous finding that ∼25% of newly synthesized Stx3 is initially targeted to the basolateral membrane in polarized MDCK cells (Sharma et al., 2006). Intriguingly, at steady - state, Stx3 is undetectable at the basolateral membrane which is consistent with our finding that Stx3 is rapidly endocytosed from this membrane. In contrast, we saw no evidence of endocytosis of Stx3 from the apical membrane. Altogether, these findings suggest that selective removal from the basolateral membrane, and stabilization at the apical membrane greatly contribute to the overall polarized distribution of Stx3 in epithelial cells.

Our results suggest that mono - ubiquitination of Stx3 acts as an internalization signal at the basolateral membrane. Similarly, mono - ubiquitination has been found to induce endocytosis of several other integral membrane proteins (Clague and Urbe, 2010; Hislop and von Zastrow, 2011; Raiborg and Stenmark, 2009; Shields and Piper, 2011). We speculate that Stx3 is recognized and ubiquitinated by an E3 ubiquitin - protein ligase that is specific to the basolateral membrane, but such an enzyme remains to be identified. A few previous studies have indicated that other syntaxins can be ubiquitinated. Stx1B was found to be poly - ubiquitinated by the novel E3 ubiquitin - protein ligase Staring which leads to its proteasomal degradation (Chin et al., 2002). Proteomic screens found evidence for ubiquitination of several syntaxins among many other proteins (Danielsen et al., 2011; Kim et al., 2011; Na et al., 2012).

We show that Stx3 is primarily mono - ubiquitinated and that this modification occurs in its polybasic juxtamembrane region. This region contains six lysine residues, all or most of which appear to be able to serve as ligation sites for ubiquitin. All other syntaxins have similar charged polybasic regions near the transmembrane domain raising the possibility that other syntaxins may be modified there, too. For example, syntaxin 5 is mono - ubiquitinated at a residue in the polybasic region as well (Huang et al., 2016). Both mono - ubiquitination events, either Stx3 or Stx5, induce dominant - negative behavior preventing cargo sorting or proper Golgi assembly, respectively. We hypothesize that ubiquitinated Stx3 may be unable to form the SNARE complex, ensuring that mistargeted Stx3 on the basolateral membrane cannot function in inappropriate membrane fusion of apically destined vesicles. This intriguing possibility remains to be investigated but would provide an elegant mechanism to enhance overall specificity in membrane trafficking.

Basolaterally endocytosed Stx3 rapidly reaches M6PR - positive endosomes (Fig. 3B). Furthermore, Stx3 is targeted to intraluminal vesicles in Rab5 - Q79L - positive endosomes (Fig. 3C). Finally, Stx3 can be recovered from apically secreted exosomes (Fig. 4). Together these results indicate that a fraction of endocytosed Stx3 is transported along the endosomal / MVB / exosomal pathway. Human urine is a relatively rich source of exosomal Stx3 (Gonzales et al., 2009) (Fig. 4) suggesting that a significant quantity of Stx3 is targeted along this route in renal tubule epithelial cells and finally excreted apically into the urinary space. The purpose of the presence of Stx3 in exosomes is unclear. One possibility is that exosomal secretion is a mechanism of eliminating excess cellular Stx3, although this would seem to be a relatively wasteful mechanism compared to lysosomal degradation. Another possibility is that Stx3 has a distinct function in exosomes although it would probably be unrelated to any membrane fusion function because topologically Stx3 protrudes into the lumen of exosomes and would be inaccessible from the outside. A third possibility – which we consider most likely - is that Stx3 functions as a trafficking adaptor directing cargo proteins into the apical exosomal secretion pathway and as a consequence it is eventually incorporated in exosomes. A finding in support of this possibility is that the ubiquitination - deficient mutant, Stx3 - 5R, appears to act as a dominant - negative inhibitor of GPRC5B into intraluminal vesicles (Fig. 3D) and its apical exosomal secretion (Fig. 4E,F). The finding that Stx3 - 5R does not affect exosomal secretion of the “generic” exosomal marker flotillin 1 (Fig. 4 D,E), and that inhibition of Stx3 expression does not affect the number of secreted exosomes (Fig. 4G,H), suggests that Stx3 functions specifically in the targeting of certain cargoes into the endosomal / MVB / exosomal pathway as opposed to be required for the generation / secretion of exosomes *per se.* Besides the abundant presence of Stx3 in urinary exosomes, GPRC5B is also secreted in urinary exosomes and its level is dramatically increased in patients with acute kidney injury (Kwon et al., 2014).

Based on the unaltered molecular weight of Stx3 recovered from exosomes (Fig. 4) we conclude that exosomal Stx3 is not ubiquitinated. However, since the non - ubiquitinatable mutant, Stx3 - 5R, fails to traffic to exosomes we conclude that ubiquitination of Stx3 is required for its entry into the endosomal / MVB / exosomal pathway, most likely at the level of endocytosis at the plasma membrane. Many membrane proteins that traffic to ILVs are ubiquitinated (Bilodeau et al., 2002; Henne et al., 2011; Raiborg et al., 2002; Shih et al., 2002), however, de - ubiquitinating enzymes typically remove ubiquitin from such proteins prior to the final incorporation into ILVs (Henne et al., 2011; Kyuuma et al., 2007; McCullough et al., 2006; Ren et al., 2008). Therefore, we propose that Stx3 is similarly de - ubiquitinated during this step but the responsible de - ubiquitinating enzyme remains to be identified.

Remarkably, the Stx3 - 5R mutant strongly inhibits the number of infectious HCMV virions secreted into the supernatant of infected cells (Fig. 5C). It is not well understood how HCMV virions are assembled and acquire their membrane envelope. However, this process occurs in an intracellular assembly site and it has been suggested that virions of HCMV and related viruses bud into a membrane organelle which subsequently fuses with the plasma membrane of the host cell to release complete, enveloped virions. Previous work has suggested that this process shares similarities with the invagination of ILVs during the formation of MVBs, and that HCMV and related viruses may exploit an exosome - like, Rab27a - dependent pathway (Calistri et al., 2007; Cepeda et al., 2010; Fraile - Ramos et al., 2010; Hurley et al., 2010; Mori et al., 2008; Pawliczek and Crump, 2009; Tandon et al., 2009).

We have previously shown that HCMV infection leads to a dramatic upregulation of Stx3 expression in the host cell and that Stx3 is required for efficient virus production (Cepeda and Fraile - Ramos, 2011). Furthermore, Stx3 is present in purified HCMV virions (Cepeda and Fraile - Ramos, 2011) suggesting that it is incorporated into the viral envelope during viral assembly. Interestingly, a virally encoded miRNA (hcmv - miR - US33 - 5p) inhibits the expression of cellular Stx3 leading to inhibition of viral replication, and it has been proposed that this mechanism facilitates establishment or maintenance of HCMV latency (Guo et al., 2015). SNAP - 23, the SNARE binding partner of Stx3 (Sharma et al., 2006), has also been shown to be required for efficient production of HCMV virions (Liu et al., 2011) suggesting that Stx3 and SNAP - 23 may function as a complex in this process. Our results confirm that knocking down the expression of endogenous Stx3 with shRNA does not interfere with the ability of HCMV to infect cells but strongly inhibits viral particle secretion (Fig. 5). Expression of shRNA - resistant Stx3 rescues this effect but expression of the shRNA - resistant Stx3 - 5R mutant does not (Fig. 5C). Furthermore, expression of Stx3 - 5R alone, over the background of endogenous wild - type Stx3, inhibits the number of secreted HCMV virions, suggesting again that this mutant acts as a dominant - negative inhibitor. Altogether, these results suggest that the ability of Stx3 to be ubiquitinated is essential for the life cycle of HCMV. We propose that ubiquitinated Stx3 functions in the assembly of enveloped HCMV virions in a trafficking pathway that is analogous to the exosomal pathway. It is possible that Stx3 may physically interact with one or more viral proteins and facilitates their trafficking to ILV - like vesicles although this remains to be investigated.

## Materials and Methods

### Reagents and antibodies

A mouse monoclonal antibody against the N - terminal residues 1 - 146 of rat Stx3 was generated. This antibody (clone 1 - 146) is reactive against the rat, mouse and human proteins and is available from MilliporeSigma (MAB2258). Anti - myc epitope tag antibody was generated from the original hybridoma cells (9E10, ATCC) and used for immunoblots and immunoprecipitation. Anti - ZO1 rat antibody (R40.76), anti - calnexin rabbit polyclonal antibody, and anti - TSG101 mouse monoclonal antibody were purchased from Santa Cruz Biotechnology. Anti - β - actin clone AC - 15 was obtained from Sigma Aldrich. Anti - myc tag, clone 4A6 from Millipore was used for immunofluorescence. Affinity - purified polyclonal antibodies against a C - terminal peptide of human SNAP - 23 have been described previously (Sharma et al., 2006). Polyclonal antibody against Munc18 - 2 (munc18b) was a kind gift from Dr. Ulrich Blank (INSERM U699, Faculté de Médecine Paris 7). Anti - Na/K ATPase mouse mAb was purchased from Affinity Bioreagents. Anti - flotillin1 mouse mAb was purchased from BD Biosciences. Secondary antibodies conjugated to DyLight 488 or 594 or and peroxidase were from Thermo Scientific and Jackson ImmunoResearch Laboratories, respectively. Secondary antibodies conjugated to IR Dye 680 or 800 were from LICOR Biosciences. Protease inhibitors, doxycycline, and nitrocellulose membranes were obtained from Sigma - Aldrich.

### Cell culture, transfection, and viruses

MDCK cells were cultured in minimal essential medium (MEM) (Corning Cellgro) containing 5% fetal bovine serum (FBS) (Omega Scientific), penicillin and streptomycin (Corning Cellgro) at 37C and 5% CO_2_. Doxycycline - inducible stable cell lines expressing Stx3 and Stx3 - 5R were made as described previously (Sharma et al., 2006). HEK293T cells were cultured in Dulbecco’s minimal essential medium (DMEM) (Corning Cellgro) containing 10% FBS, penicillin and streptomycin at 37C and 5% CO_2_. Transient transfections were carried out using Lipofectamine 2000 (Life Technologies) or TurboFect (ThermoFisher) per manufacturer’s instruction.

Immortalized human foreskin fibroblast (BJ1) cells were from Clontech and human melanoma MeWo cells were a gift from Dr Luís Montoliu (CNB, Madrid, Spain). Cells were maintained as recommended by suppliers. MeWo and BJ1 transduced cells were selected in media containing 10 and 2 μg/mL puromycin respectively. BJ1 cells were transduced with lentiviruses expressing a c - myc - Stx3 - wt or a c - myc - Stx3 - 5R mutant construct resistant to shRNA inhibition. BJ1 shRNA - resistant c - myc - Stx3 - wt and - 5R expressing cells were sorted with an ALTRA HyPerSort flow cytometer (Beckman Coulter, Inc., Palo Alto, USA).

A recombinant strain of HCMV AD169 expressing GFP under the control of the HCMV early promoter beta 2.7 gene that is expressed from 8 hpi, RecCMV (McSharry et al., 2001), was propagated on BJ1 cells and titrated as previously described (Cepeda et al., 2010).

Lentiviral vectors for shRNA - mediated gene silencing were prepared with pMDG, p8.91 and retroviral expression plasmids encoding scrambled control (SHC002) and Stx3 shRNA TRCN0000065016 (#304) (Mission^®^ TRC - Hs shRNA libraries, Sigma Aldrich) (Moffat et al., 2006) as described (Naldini et al., 1996).

### Plasmids and shRNA

pcDNA4 - Stx3 expression constructs were described previously (Sharma et al., 2006). QuickChange Site - directed mutagenesis kit (Agilent Technologies) was used to generate mutations in the pcDNA4 - Stx3 expression construct per manufacturers instructions. pGFP - C2 - hRab5A and pGFP - C2 - hRab5A - Q79L were gifts from Dzwokai Ma (University of California, Santa Barbara). pEGFP - GPRC5B plasmid was a gift from Keith Mostov and Sang - Ho Kwon (University of California, San Francisco). mCherry - Rab5CA(Q79L) was a gift from Sergio Grinstein (Addgene plasmid # 35138) (Bohdanowicz et al., 2012).

shRNA - resistant c - myc - Stx3 (GenScript, New Jersey, USA) and c - myc - Stx3 - 5R constructs were extended with Gateway^®^ recombination sequences and transferred via pDONR201 (Invitrogen) to the lentiviral plasmid pLenti - PGK - Neo - DEST(w531 - 1) (Campeau et al., 2009) (obtained through Addgene, Massachusetts, USA). Recombinant lentiviruses were prepared as described above.

### Co - immunoprecipitation

Polarized MDCK cells were lysed in a buffer containing 50mM Hepes - KOH pH 7.4, 50mM potassium acetate, 1% Triton X - 100, and 100μM PMSF for 30 minutes rotating at 4°C. Samples were centrifuged at 10,000g for 10 minutes at 4°C. Resulting supernatant was pre - cleared with CL2B sepharose beads for 20 minutes at 4°C. Pre - cleared supernatants were incubated with protein - A beads cross - linked to anti - myc mouse IgG antibody (GE Healthcare Life Sciences) overnight rotating at 4°C. Beads were washed three times with lysis buffer followed by one wash with lysis buffer minus Triton X - 100. Beads were resuspended in sample buffer and subjected to SDS - PAGE and Western blotting.

### Immunofluorescence microscopy

Cells were fixed with 4% paraformaldehyde, and treated with permeabilization/blocking buffer (PBS containing 5% donkey serum and 0.2% Triton X - 100). Primary and secondary antibodies were diluted in permeabilization / blocking buffer. Cells were stained with DAPI (Sigma - Aldrich) and mounted with ProLong Gold mounting medium (Life Technologies).

For endocytosis assays, MDCK cells were cultured on Transwell membrane filters (Corning Cellgro) with complete media in the basolateral chamber and serum - free media in the apical chamber for 4 days to ensure polarization. Anti - myc in supplemented MEM (with 20mM Hepes, pH 7.4 and 0.6% BSA) was added either to the basolateral or to the apical chamber for 20min at 4°C. Cells were washed three times with ice - cold supplemented MEM before being incubated at 37°C for the indicated times and processed as above. Images were acquired using an Oympus Fluoview FV1000S Spectral Laser Scanning Confocal microscope using an Olympus UPLFLN 60× oil - immersion objective. FIJI software was used to generate plot - profile graphs for colocalization analysis. Composite images were assembled in Adobe Photoshop.

### Exosome purification and quantification

Human urine or media from cell cultures were subjected to three centrifugation steps; 1,000g for 10 minutes, 10,000g for 10 minutes, and 100,000g for 60 minutes. At each step, the supernatant was transferred to a new tube. The resulting 100,000g pellet was resuspended in Laemmli buffer and subjected to SDS - PAGE and Western blotting. Bands were quantified with Odyssey Software (LiCOR), bar graphs and statistics were performed in Prism 7 (GraphPad).

For exosome quantification, 1 x 106 MeWo cells transduced with scrambled control and Stx3 shRNAs growing in 60 - mm tissue culture plates were incubated in 2 mL of tissue culture medium containing 1% FCS. After 48 hours, the cells were lysed to analyse the levels of Stx3 and tubulin by immunoblot, and the supernatants were harvested to determine the number of exosomes and their size distribution by Nanoparticle Tracking Analysis (Carr, 2007). Briefly, cell culture supernatants were centrifuged at low - speed in sequential steps, and then clarified to eliminate cell debris. Clarified supernatants were diluted (1/5) in HBSS and analysed with the use of NanoSight LM10 instrument (NanoSight Ltd., UK) as described (Dragovic et al., 2011). NanoSight was calibrated with 100 nm and 400 nm fluorescent calibration beads (Malvern, UK).

### HCMV infection in BJ1 cells

BJ1 parental cells and cells expressing an shRNA - resistant c - myc - Stx3 - wt or - 5R construct plated at 0.8 x 105 cells in 16 mm wells in 24 - well plates were transduced with scrambled control or Stx3 shRNAs. At 2 days post transduction, cells were infected with RecCMV at an MOI of 0.3 and 4 hours later puromycin was added to the culture to select transduced cells. At 4 days post infection (dpi), a fraction of the cells were fixed, analyzed by FACS and the number of infected cells was quantified by GFP expression, another fraction of the cells were lysed to quantify the expression of STX3 by Western blot, whereas the remainder of the cells were washed and fresh medium was added to collect viruses secreted into the supernatants. At 5 dpi supernatants were harvested and extracellular infectious particles were titrated on fresh BJ1 cells as previously described (Fraile - Ramos et al., 2007). Cells were fixed at 56 hours post infection, analysed by flow cytometry and the number of infectious cells, i.e. number of infectious virus particles, was assayed by GFP expression.

## Acknowledgments

We thank Dzwokai Ma, Keith Mostov, Sang - Ho Kwon, Sergio Grinstein, Ulrich Blank and Luís Montoliu for gifts of reagents and advice. We specially thank Raquel Bello - Morales for help with the Nanoparticle Tracking Analysis. We acknowledge the use of the UCSB NRI - MCDB Microscopy Facility and the Spectral Laser Scanning Confocal supported by the Office of The Director, National Institutes of Health under Award # S10OD010610. This work was supported by grants from the NIH (DK62338), and the California Cancer Research Coordinating Committee to T.W., by a grant BFU2012 - 35067 from the Spanish Ministry of Economy and Competitiveness (MINECO) to A.F. - R., and a Postdoctoral Fellowship from the Spanish Ministry of Education and Science to E.R.

